# Novel ACE2-IgG1 fusions with improved *in vitro* and *in vivo* activity against SARS-CoV2

**DOI:** 10.1101/2020.06.15.152157

**Authors:** Naoki Iwanaga, Laura Cooper, Lijun Rong, Brandon Beddingfield, Jackelyn Crabtree, Ralph A. Tripp, Xuebin Qin, Jay K. Kolls

## Abstract

SARS-CoV2, the etiologic agent of COVID-19, uses ACE2 as a cell entry receptor. Soluble ACE2 has been shown to have neutralizing antiviral activity but has a short half-life and no active transport mechanism from the circulation into the alveolar spaces of the lung. To overcome this, we constructed an ACE2-human IgG1 fusion protein with mutations in the catalytic domain of ACE2. This fusion protein contained a LALA mutation that abrogates Fcrγ binding, but retains FcRN binding to prolong the half-life, as well as achieve therapeutic concentrations in the lung lavage. Interestingly, a mutation in the catalytic domain of ACE2, MDR504, completely abrogated catalytic activity, but significantly increased binding to SARS-CoV2 spike protein *in vitro*. This feature correlated with more potent viral neutralization in a plaque assay. Parental administration of the protein showed stable serum concentrations with a serum half-life of ∼ 145 hours with excellent bioavailability in the epithelial lining fluid of the lung. Prophylactic administration of MDR504 significantly attenuated SARS-CoV2 infection in a murine model. These data support that the MDR504 hACE2-Fc is an excellent candidate for pre or post-exposure prophylaxis or treatment of COVID-19.

## Introduction

SARS-CoV-2 is the etiologic agent of the current COVID-19 pandemic. Similar to SARS-CoV and other betacoronaviruses it uses the angiotensin-converting enzyme 2 (ACE2) receptor [1]. ACE2 is expressed in the nasal respiratory epithelium, in the conducting airway in type II pneumocytes [2] and elsewhere [3]. It has been hypothesized that the spike protein of SARS-CoV-2 has stronger binding to human ACE2 compared to SARS-CoV [4, 5]. Recent surface plasmon resonance assays with soluble ACE2 binding SARS-COV-2 spike protein had a binding affinity in the nM to pM range [6]. Thus, the soluble version of ACE2 may have potent viral neutralization activity. ACE2 is a type I transmembrane domain with a 740 amino acid ectodomain [7]. Soluble ACE2 has been shown to bind to SARS-CoV and SARS-CoV-2 spike proteins and block viral entry [8]. Soluble ACE2 has been administered to humans with pulmonary hypertension [9] and acute respiratory distress syndrome (ARDS) [10] at dose ranges of 0.1 to 0.8 mg/kg and shown to be well tolerated. However, the pharmacokinetic/pharmacodynamic (PK/PD) of soluble ACE2 is not ideal for sustained viral neutralization *in vivo* and is not engineered to be transported from the circulation into the epithelial lining fluid of the lung. ACE2-IgG Fc fusion proteins have been reported to also bind virus and neutralize SARS-CoV-2 pseudoviruses *in vitro* [6]. Moreover, several mutants in the catalytic domain have been reported that also bind and neutralize virus [6]. However, ACE2-IgG Fc fusion proteins retain FcRγ binding which may compromise serum stability or activate FcRγ on myeloid cells, which may be problematic in COVID-19.

Thus, we engineered four distinct ACE2-Fc IgG1 fusions all with the LALA mutation to abrogate FcRγ binding but retain binding to the neonatal Fc receptor which is important for serum stability [11] as well as transport into the lung [12]. In addition to the wild-type ACE2 ectodomain, we engineered several mutations to inactivate catalysis of angiotensin such as MDR504 and MDR505. Constructs included an IgGκ leader sequence and an IEGR linker between ACE2 and the CH2 and CH3 domains of human IgG1. All constructs expressed proteins consistent with homodimers after transfection in HEK293 cells. All constructs bound monomeric SARS-CoV-2 receptor binding domain as well as trimeric spike protein [13]. However, the binding of the MDR504 mutant was superior at both room temperature and 37 °C. This correlated with enhanced neutralization of virus in a Vero E6 cell plaque assay. Furthermore, the MDR504 mutant had similar serum stability as wild-type ACE2-Fc but appears to have higher levels in the epithelial lining fluid of the lung after parenteral administration to C57Bl/6 mice. Based on these studies, the MDR504 mutant appears to be an excellent candidate to move forward in terms of preventing or treating COVID-19.

## Methods

### Mice

Female wild type C57BL/6J mice 6-10-week-old were used for *in vivo* studies. The mice were bred in-house or purchased from The Jackson Laboratory. All experiments were performed using sex- and age-matched controls and approved by the Institutional Animal Care and Use Committee of Tulane University.

### Generation of different constructs of human ACE2 fusion proteins

The DNA sequences of the extracellular domains of *ACE2* and IgG1 were synthesized by Genscript and cloned into pcDNA3.1. Transient transfection was performed using Lipofectamine™ 3000 Transfection Reagent (Invitrogen) in 293T cells. The collected supernatants were collected and purified by protein G-sepharose (Thermo Fisher). The concentration and purity were confirmed by measuring the UV absorbance at wavelength of 280 nm, BCA assay (Thermo Fisher) and human IgG ELISA (Thermo Fisher).

### Western blotting

Following the removal of the supernatants, cell lysates were dissolved in PBS (50 mg/ml) containing protease inhibitors (Thermo Scientific) and 1 mM PMSF. BCA assay was performed to quantify protein and 5.0 μg protein was used for Western blotting. Western blots were performed using 7.5% SDS-PAGE gels (Bio-Rad) under the non-reducing or reducing condition with 2.5% 2-mercaptoethanol and transferred to PVDF membranes. The blot was probed with goat anti-human IgG-HRP (Southern Biotech). After incubation with IgG-HRP-conjugated anti-human antibody, membranes were washed and incubated with SuperSignal West Pico Chemiluminescent Substrate (Thermo Scientific). Signal was detected using Bio-Rad ChemiDoc MP imaging system.

### ELISA for human ACE2 and spike protein

ELISA plates were coated with 2 μg/mL recombinant spike glycoprotein receptor binding domain (RBD) from SARS-CoV-2 (BEI RESOURCES) or recombinant S1 subunit (RayBiotech) overnight at 4 °C. Coated plates were washed with washing buffer (0.05% Tween 20 in PBS), blocked for 2 h at room temperature with blocking buffer (1% BSA and 0.1% Tween 20 in PBS), and washed before the addition of the supernatants or cell lysates from transfected 293T cells. After 2 h incubation at RT, or 1 h incubation at 37 °C, the plates were washed and incubated with goat anti-human IgG conjugated with horseradish peroxidase (Southern Biotech) diluted 1/5,000 in assay diluent (0.5% BSA and 0.05% Tween 20 in PBS) for 1 h at RT, or for 30 min at 37 °C, TMB peroxidase substrate (Southern Biotech) was added to each well. Absorbance was read at 450 nm on a microplate reader (BioTek).

### Pseudovirus Production

Pseudoviruses for antibody screening were generated using the following plasmids: S-Tag of SARS (pcDNA3.1(+)-SARS-S), S-Tag of SARS2 (pcDNA3.1(+)-SARS2-S), vesicular stomatitis virus (pcDNA3.1(+)-VSV-G), and the HIV-1 pro-viral vector pNL4-3.Luc.R^-^E^-^ which were obtained through the NIH AIDS Research and Reference Reagent Program. All pseudoviruses were produced by transient co-transfection of 293T cells using a polyethyleneimine (PEI)-based transfection protocol. Five hours after transfection, cells were washed with phosphate-buffered saline (PBS), and 20 mL of fresh media was added to each 150 mm plate. Twenty-four hours post transfection, the supernatant was collected and filtered through a 0.45 µM pore size filter and stored at 4 °C prior to use.

### Antibody neutralization Assay

Targeted 293T cells were transfected with pcDNA3.1(+)-humanACE2 and pCSDest-TMPRSS2 plasmids for 6 h. The cells were then trypsinized and seeded 1×10^5^ cells/well in DMEM complete into 96-well plates (100 μL/well) then incubated for 16 hours at 37 °C and 5% CO_2_. SARS-1, SARS-2, and VSV pseudoviruses were incubated with the test samples at room temperature for 1 h, and then added to the target cells in 96-well plates. Plates were incubated for 48 hours and levels of viral infection were determined by luminescence using the neolite reporter gene assay system (PerkinElmer). Virus alone was used as a control and data was normalized to the control.

### Human ACE2 neutralization of SARS-CoV2 by plaque assay

Vero E6 cells were plated in a 6 well plate at 8 × 10^5^ cells per well and incubated overnight. Each construct of hACE2-Fc was preincubated with SARS-CoV2 virus for 10 minutes before infection in 1 ml media. The cells were washed once with PBS and infected at a MOI of 0.01 with the compounds for 1 h. Following infection, the supernatants containing the virus and compounds were removed and 3 ml overlay media containing each compound were added to the wells and incubated for additional 4 days. Post infection, the cells were fixed and stained to visualize plaques.

### Pharmacokinetics study

Mice were injected with 4 mg / kg body weight MDR504 via retro-orbital (i.v.) and then the serums and bronchoalveolar lavage fluid were collected at each time point. The concentration of MDR504 were analyzed by detecting IgGFc by ELISA. We used purified anti-human IgG Fc antibody (Biolegend) as a capture antibody and anti-Human IgG Fc, Multi-Species SP ads-HRP (SouthrenBiotech) as a detection antibody and the other procedure is same as mentioned above.

### In vivo evaluation of MDR504

All animals are cared for in accordance with the NIH guide to Laboratory Animal Care. The Institutional Biosafety Committee approved the procedures for sample handling, inactivation, and removal from a BSL3 containment. For this study, we utilized the murine model of SARS-CoV2 were C57Bl/6 mice were first inoculated with adenovirus encoding human ACE2 (Vector Biolabs). Four days later, mice received 5 × 10^4^-2 × 10^5^ pfu of SARS-CoV-2 intranasally. To evaluate MDR504, mice were dosed with wild-type hACE2-Fc or MDR504 three to four hours prior to SARS-CoV2 infection. Mice were euthanized 72 hours post infection, tissue samples were collected in Zinc formalin (Anatech), embedded in paraffin and 5 um thick sections were cut, adhered to charged glass slides. 5um sections of Zinc Formalin-fixed, paraffin-embedded lung were mounted on charged glass slides, baked overnight at 56oC and passed through Xylene, graded ethanol, and double distilled water to remove paraffin and rehydrate tissue sections. A microwave was used for heat induced epitope retrieval. Slides were heated in a high pH solution (Vector Labs H-3301), rinsed in hot water and transferred to a heated low pH solution (Vector Labs H-3300) where they were allowed to cool to room temperature. Sections were washed in a solution of phosphate-buffered saline and fish gelatin (PBS-FSG) and transferred to a humidified chamber. Tissues were blocked with 10% normal goat serum (NGS) for 40 minutes, followed by a 60 minute incubation with polyclonal guinea pig anti-SARS-CoV1 antibodies (1:1000) (NR-10361, BEI Resoruces). Slides were transferred to the humidified chamber and incubated, for 40 minutes, with secondary antibodies tagged with various Fluor fluorochromes and diluted to a working concentration of 2 ug/mL. Slides were mounted using a homemade anti-quenching mounting media containing Mowiol (Calbiochem #475904) and DABCO (Sigma #D2522) and imaged with a Zeiss Axio Slide Scanner.

### Statistical analysis

Statistical analysis was performed with GraphPad Prism 8.0. P values < 0.05 was evaluated statistically significant. Comparisons between two normally distributed groups were performed by simple 2-tailed unpaired student’s t-test. For multiple groups comparisons, we used one-way or two-way ANOVA with Tukey’s post-hoc analysis. Values are represented as means ± SEM. P values are annotated as follows (*) ≤0.05, (**) ≤0.01, (***) ≤0.001, and (****) ≤0.0001.

## Results

### Expression of hACE2-Fc constructs

The amino acid sequences are listed in Supplemental Table I. After transient transfection of the constructs into 293T cells, human IgG was readily detected by ELISA (data not shown). Proteins were run on a reduced SDS-PAGE gel and migrated at ∼ 140 kDa consistent with the predicted molecular weight of the monomer (Supplementary Figure 1A). SDS-PAGE analysis in non-reducing conditions revealed migration consistent with a dimeric protein (Supplementary Figure 1B).

### SARS-CoV2 RBD and Spike Binding

Proteins were purified from the supernatant of 293T cells, and we assayed binding to immobilized RBD or spike by ELISA. In experiments run at room temperature, we observed higher binding of the MDR504 and MDR505 hACE2-Fc and a lesser extent to the MDR503 hACE2-Fc compared to wild-type ACE2-Fc (Figure 1A and B). This increase in binding was more dramatic when assayed at 37°C (Figure 1C and D) where binding of the MDR504 mutant was superior for the both RBD assay as well against the trimeric spike protein. To verify that the MDR504 mutant lacked ACE2 catalytic activity we performed an *in vitro* catalysis assay using a fluorogenic peptide that contains an ACE2 cleavage site. WT ACE2 showed substantial catalysis of the peptide whereas the MDR504 mutant completely lacked any detectable activity (Figure 1E).

**Figure 1.**
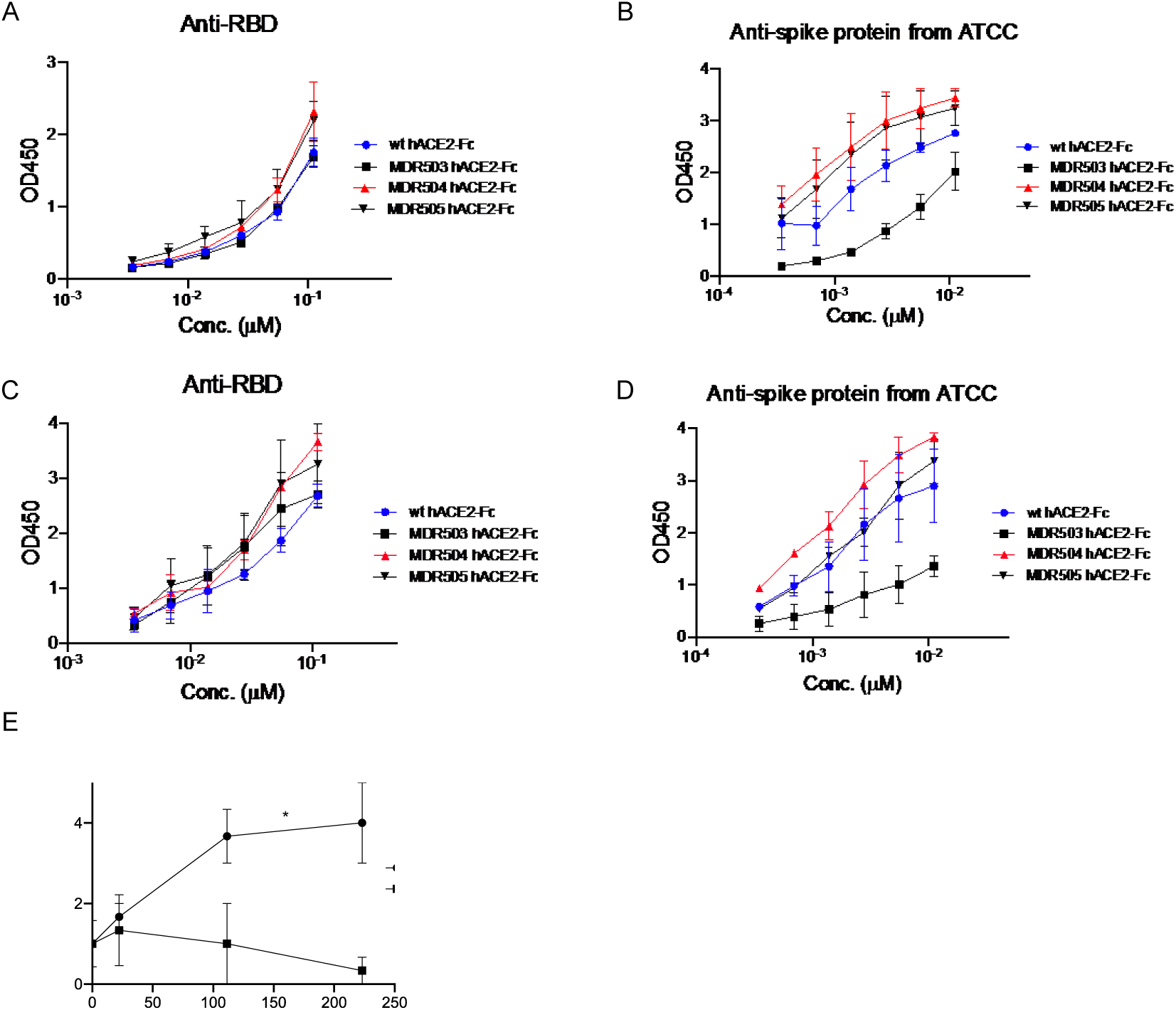
In vitro biding of MDR504. Binding of RBD (A, C) or spike protein (B, D) and hACE2 by ELISA. Assays were performed at room temperature (A,B) or at 37°C (C, D). (E) Catalytic activity of WT or the MDR504 mutant in a fluorogenic ACE2 assay (*n* = 3). Significant differences are designated using one-way ANOVA followed by Tukey’s multiple comparisons test (B, D) and unpaired t-test (E) based on the AUC. *, P < 0.05; **, P < 0.01; ***, P < 0.001.

### Neutralization of SARS-CoV2 infection

Using a pseudotyped model of SARS-CoV-2 we found superior neutralization with the MDR504 mutant and the MDR505 double mutant compared to wild type human ACE2 Fc IgG1 (Figure 2A, 2B). Next, we examined neutralization of SARS-CoV-2 using a plaque assay in Vero E6 cells. Initial studies were done at 50 μg/ml (223 nM) based on studies with Pavalizumab, an anti-RSV monoclonal antibody, showed that effective anti-RSV trough concentrations *in vivo* were ∼ 40 μg/ml [14]. 50 μg/ml completely neutralized SARS-CoV-2 *in vitro* (Figure 2C). Similar, but slightly less activity was observed with the MDR503 mutant (Figure 2C). Next, we compared the wild-type protein to the dual mutant MDR505 and the MDR504 mutant at the lower concentration of 1 and 10 μg/ml. Here, both mutants neutralized the virus and considerably improved over the wild-type protein (Figure 2D). Moreover, we confirmed the significant improvement in the IC_50_ of the MDR504 mutant (Figure 2E).

**Figure 2.**
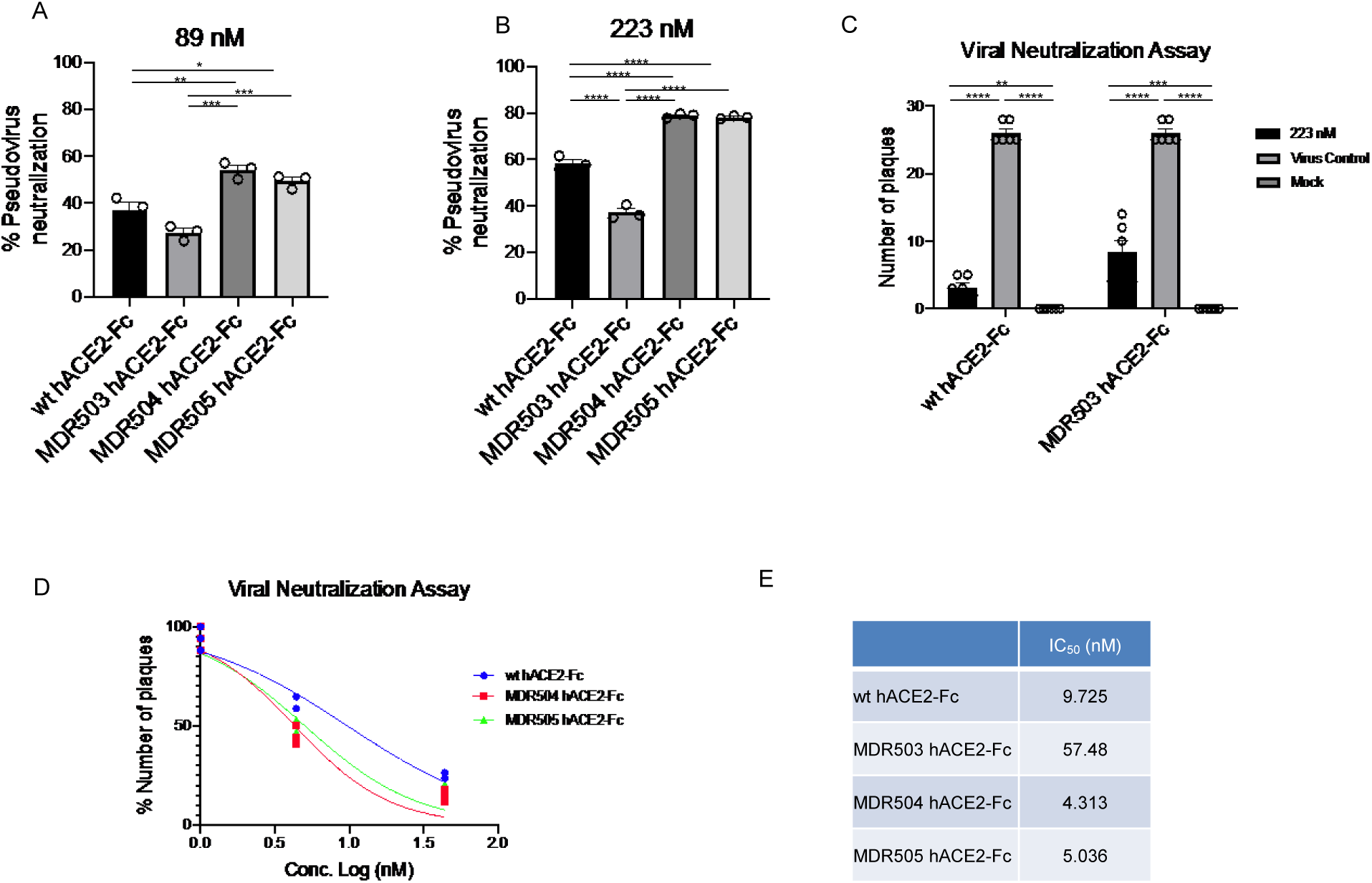
In vitro efficacy of MDR504. Neutralization of pseudovirus were compared among the different constructs in (A) 89 nM and (B) 223 nM of various hACE2-IgG1 fusions. MDR504 mutant and MDR505 double mutant showed significantly higher neutralization. Similarly, SARS-CoV2 neutralization with WT and MDR503 mutant at 223 nM attested the result by plaque assay (C). IC_50_ of the different ACE2-IgG1 fusions in the placque assay (D). Significant differences are designated using one-way ANOVA (A, B, C) and two-way ANOVA (D) followed by Tukey’s multiple comparisons test. *, P < 0.05; **, P < 0.01; ***, P < 0.001; ****, P < 0.0001. (E) Calculated IC_50_ of each construct based on the placque assay data.

### Serum Stability and BAL Fluid Concentrations

C57BL/6 mice were injected with wild-type ACE2-Fc or the MDR504 mutant and serum was collected at 0, 1, 24, or 72 h. Mice were euthanized at 6h and 72 h and underwent bronchoalveolar lavage (BAL) to measure ACE2-Fc in the lung. The MDR504 mutant had a slightly higher peak concentration in serum and a half-life of approximately 145 h (Figure 3A0. The MDR504 mutant had slightly higher concentrations in BALF at 72 h compared to the wild-type protein (Figure 3B).

**Figure 3.**
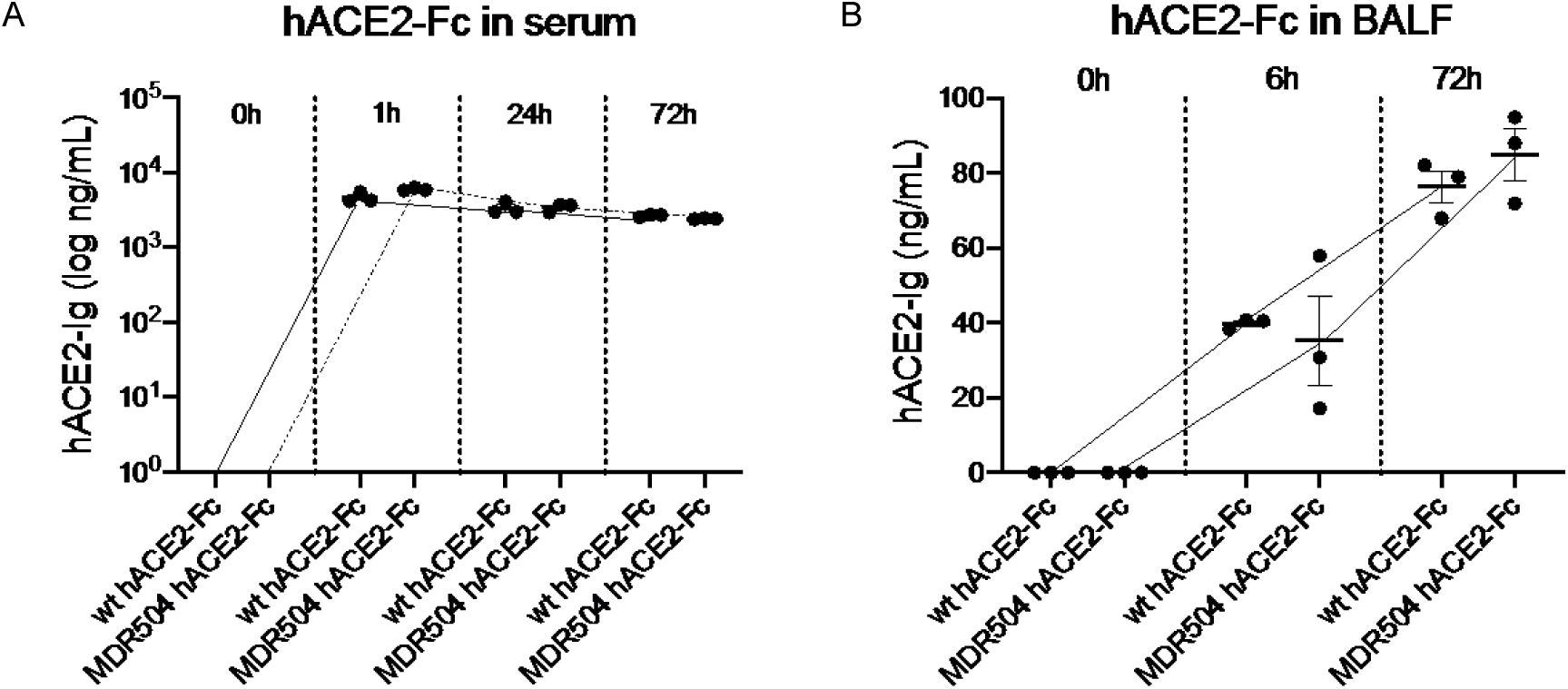
In vivo PK of MDR504. In vivo pharmacokinetics of the WT and MDR504 mutant ACE2-IgG1 was assayed after intravenous injection of 4 mg / kg body weight of protein assayed in in serum (A) and BAL fluid (B).

**Figure 4.**
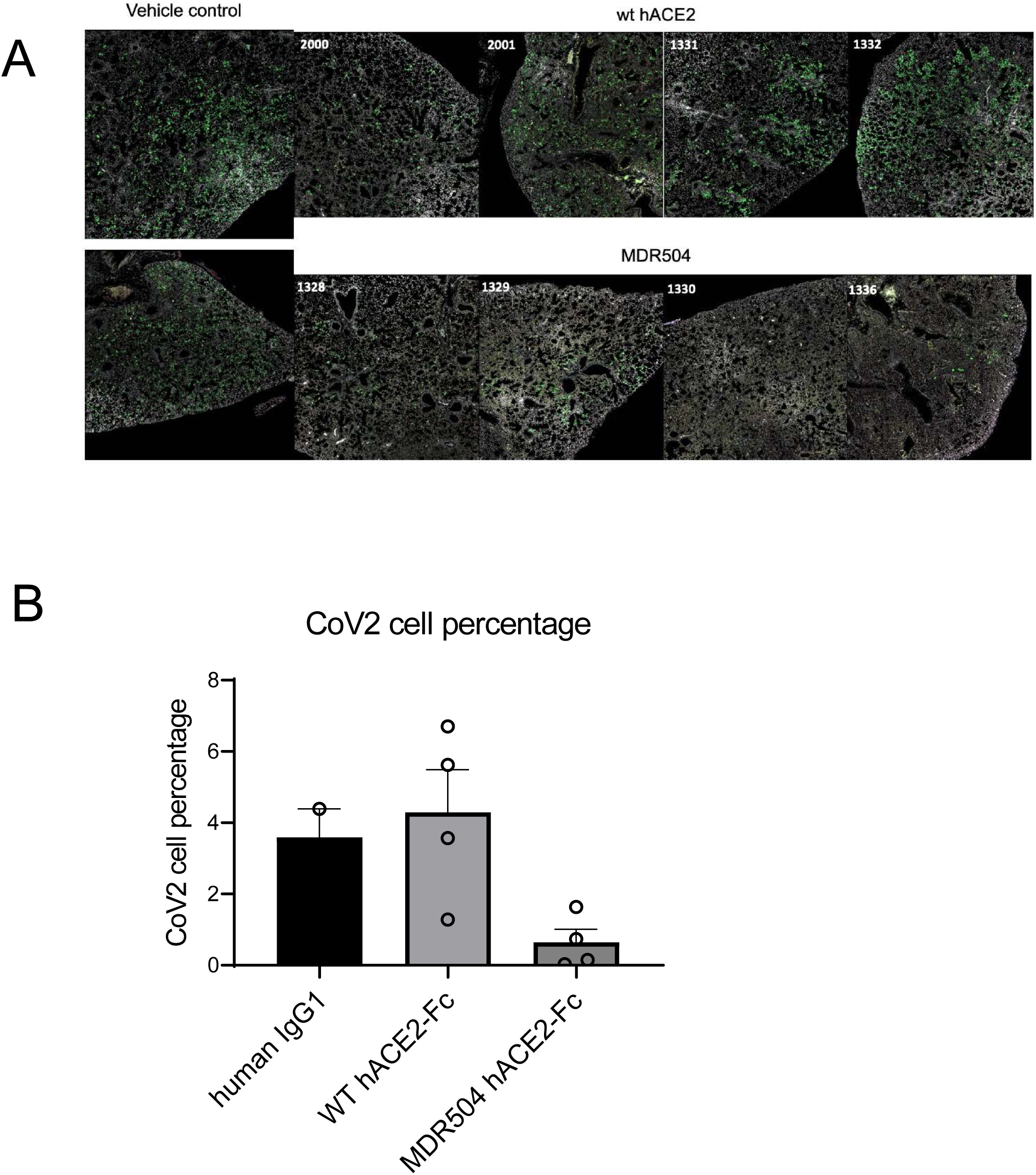
In vivo efficacy of MDR504. C57Bl/6 mice were administered AdhACE2. 4 days later mice were treated with vehicle, or 15 mg/kg WT-ACE2 or mutant ACE2-IgG1 IV. 4 hours later mice were infected intranasally with SARS-Cov2. Mice were euthanized three days post SARS-CoV2 inoculation in ABSL3. SARS-CoV2 infection was assayed by IFC.

### In vivo efficacy experiments

Histological analysis of vehicle treated mice showed widespread SARS-COV2 infection in the distal lung with approximately 4% of the cells infected (Figure X). This was largely unaffected by wild-type ACE2-Fc prophylaxis. In contrast, MDR504 had significantly less infected cells with only 0-1% of cells positive for SARS-CoV2 staining (Figure X).

## Discussion

One mechanism that has made SARS-CoV-2 more infectious than the 2002 SARS-CoV epidemic is potentially the higher affinity of SARS-CoV-2 spike protein to the human ACE2 [4, 5]. However, this increased affinity represents a potential therapeutic target to block viral entry. To this end, we made ACE2-Fc fusions with mutations in the ACE2 catalytic domain as well as in the IgG1 constant region to abrogate FcRγ binding. Interestingly, we found that the MDR504 mutation had greater binding the SARS-CoV-2 RBD and spike protein. This translated to a lower IC_50_ value for infectious viral neutralization and in a pseudovirus assay. At this time, the structural basis for this remains unclear. Cryo-EM studies of SARS-CoV-2 RBD has been shown to bind the NH2 terminus of human ACE2 [15]. However, the RBD also binds to residues K353, G354, and D355 [15] and thus it is possible that the MDR504 mutation affects this binding. The other reported ACE2-Fc mutants showed equivalent binding and pseudovirus neutralization [6]. The authors of this paper made H374N and H378N mutations that have putatively reduced ACE2 catalytic activity but that was not specifically assayed as part of their study [6]. However, our IC_50_ data is in agreement with their calculated *K*_*d*_ using surface plasmon resonance where the *K*_*d*_ was reported to be 11.2 nM [6]. Thus, the MDR504 mutation may have a structural basis for enhanced neutralization over wild-type ACE2 and that will be the subject of future research. Importantly our constructs also showed activity against infectious SARS-CoV-2. The MDR504 mutant also showed excellent stability in serum and achieved therapeutic levels in bronchoalveolar lavage (BAL) fluid in a murine PK/PD study.

Taken together, this reagent may be useful as pre- or post-exposure prophylaxis or as therapy for COVID-19. This technology may also complement vaccine technology, as it may be useful in subjects that may not be good candidates for vaccines such as patents with hematologic or other malignancies, or those that are undergoing immunosuppressive therapy for organ transplantation or autoimmune disease.

## Acknowledgements

This work was supported by the following NIH grant R35HL139930 (JK) and R21OD024931 (XQ). The following reagent was produced under HHSN272201400008C and obtained through BEI Resources, NIAID, NIH: Spike Glycoprotein Receptor Binding Domain (RBD) from SARS-Related Coronavirus 2, Wuhan-Hu-1, Recombinant from HEK293 Cells, NR-52306. We are grateful for technical help from Cecily C Midkiff and Dr. Robert Blair in the histological core of Tulane National Primate Research Center and Christopher J Monjure in the Tulane ABSL3 core.

---

**Supplementary Figure 1.**
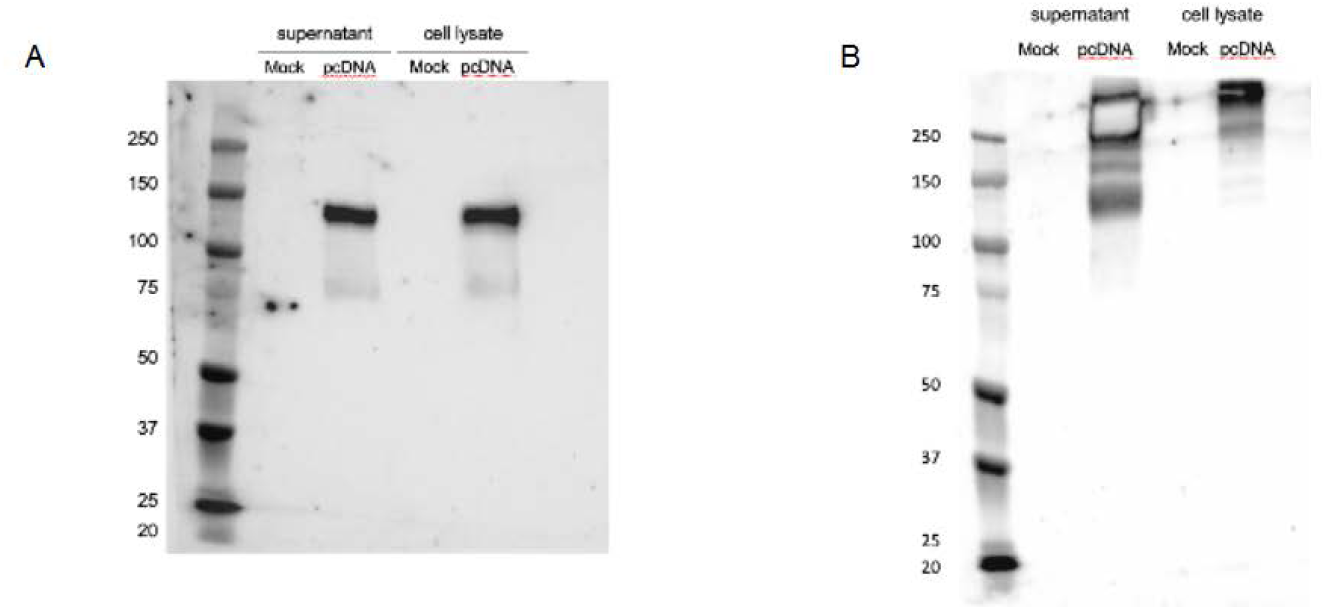
(A) Western blotting under reducing condition with 2.5% 2-mercaptoethanol of hACE2 expression in cell supernatants of transfected cells as well as cell lysates. (B) Western blotting of hACE2-IgG1 without reducing conditions.

